# Dopaminergic but not Cholinergic Neurodegeneration is Correlated with Gait Disturbances in PINK1 Knockout Rats

**DOI:** 10.1101/2021.04.15.440072

**Authors:** VM DeAngelo, JD Hilliard, GC McConnell

**Affiliations:** Department of Biomedical Engineering, Stevens Institute of Technology, Hoboken, NJ 07030; Department of Neurosurgery, University of Kentucky, Lexington, KY 40536

**Keywords:** DA = dopamine, P1KO = PTEN-induced putative kinase 1 knockout, PBS = phosphate buffered saline, PD = Parkinson’s disease, PPN = pedunculopontine nucleus, SNc = substantia nigra pars compacta, VTA = ventral tegmental area, WT = wild-type

## Abstract

Parkinson’s disease (PD) is a progressive neurodegenerative disorder characterized by gait dysfunction in later stages of the disease. PD hallmarks include a decrease in stride length, run speed, and swing time; an increase in stride time, stance time, and base of support; dopaminergic degeneration in the basal ganglia; and cholinergic degeneration in the pedunculopontine nucleus (PPN). A progressive animal model of PD is needed to identify treatments for gait dysfunction. The goal of this study was to quantify progressive gait degeneration in PTEN-induced putative kinase 1 knockout (P1KO) rats and investigate neurodegeneration as potential underlying mechanisms. Gait analysis was performed in male P1KO and wild-type rats at 5 and 8 months of age and immunohistochemical analysis at 8 months. Multiple parameters of volitional gait were measured using a runway system. P1KO rats exhibited significant gait deficits at 5 months, but not 8 months. Gait abnormalities improved over time suggesting compensation during behavioral testing. At 8 months a 15% loss of tyrosine hydroxylase (TH) in the striatum, a 27% loss of TH-positive cells in the substantia nigra pars compacta, and no significant loss of choline acetyltransferase-positive cells in the PPN was found. Dopaminergic cell loss may contribute to gait deficits in the P1KO model, but not cholinergic cell loss. The P1KO rat with the greatest dopamine loss exhibited the most pronounced PD-like gait deficits, highlighting variability within the model. Further analysis is required to determine the suitability of the P1KO rat as a progressive model of gait abnormalities in PD.

## Introduction

Parkinson’s disease (PD) is a neurodegenerative movement disorder characterized by the progressive loss of dopaminergic neurons in the substantia nigra pars compacta (SNc) that project to the striatum [1,2,3] as well as degeneration of other motor nuclei, such as loss of cholinergic neurons in the pedunculopontine nucleus (PPN) [4]. Cardinal symptoms of PD include resting tremor, rigidity, bradykinesia, postural instability, and gait disturbances. Gait disturbances are among the most common and debilitating symptoms of advanced PD (>40% dopamine (DA) loss) [1]. Symptoms present as shuffled gait, decreased stride length and velocity, and increased base of support [5] associated with increased falls [6].

Animal models of PD enable the investigation of the neuropathophysiology and the development of novel treatments. There are multiple models available that exhibit one or more features of clinical PD, however, no current rodent model exhibits gait disturbances with a timeline similar to the natural progression of the disease [7]. An ideal model of PD should be age dependent and progressive [8]. The most widely used rodent model of PD is the 6-hydroxydopamine (6-OHDA) hemiparkinsonian model. Administration of 6-OHDA causes degeneration of the dopaminergic neurons in the SNc, resulting in gait deficits as early as 4 days post-lesion [9,10]. The lesion is restricted to one hemisphere of the brain, causing an acute asymmetrical presentation of symptoms, unlike typical PD.

A possible alternative to the 6-OHDA model is the PINK1 (PTEN-induced putative kinase 1, (P1)) knockout (KO) in which the PINK1 gene is deleted. Loss of function mutations in P1, important for preserving mitochondrial function in the brain, is linked to early onset PD with slow progression [11,12]. The P1KO rat is a commercially available genetic rodent model initially shown to experience 50% dopamine depletion in the SNc and an increase of dopamine in the striatum at 8 months of age, compared to healthy wild-type (WT) rats. P1KO rats exhibit locomotor deficits beginning as early as 4 months of age including decreased hindlimb grip strength, increased number of foot slips on a tapered balance beam, and abnormal gait [13]. It is unclear if these locomotor deficits occur in tandem with abnormal gait displayed as rats walk in a straight line. Prior studies reported motor abnormalities, but measurements of DA loss within the SNc and striatum have been inconsistent. Dopaminergic degeneration in the P1KO model was reported to range from no loss [14,15] to significant loss [16,17], however, behavioral deficits were reported by all. These results indicate that the neural mechanism for gait dysfunction in the P1KO model may be due to neurodegeneration of other brain regions and networks, such as the PPN.

The PPN, located in the dorsal tegmentum of the midbrain and upper pons, is an understudied brain region in rodents. Post-mortem studies of PD patients demonstrated cholinergic neuronal loss in the PPN of up to 50% [4,18,19]. Loss of cholinergic neurons in the PPN is believed to synergistically exacerbate neuronal loss in the SNc resulting in gait and balance deficits in patients. Pienaar *et al*. [4] analyzed mitochondrial respiratory chain subunit expression patterns in remaining PPN neurons in PD and found a significant decrease in mitochondrial protein expression, indicative of mitochondrial injury. We hypothesized that gait dysfunction in P1KO rats is attributed to cholinergic loss in the PPN in light of the inconsistency observed for DA loss in the SNc, and the mitochondrial injury observed in P1 mutations and PPN neurons in PD.

In this study we investigated the suitability of P1KO rats as a progressive model of gait dysfunction in PD. We measured voluntary gait in P1KO rats and age-matched WT control rats at 5 months and 8 months of age. Further, we quantified dopaminergic degeneration in the SNc, striatum and VTA, and cholinergic degeneration in the PPN to understand the mechanism underlying gait disturbances.

## Material and Methods

### Animals and housing

Adult male Long-Evans hooded P1KO rats (*n* = 4) and Long-Evans hooded WT rats (*n* = 4) derived from the breeding of the P1KO rats were purchased from Horizon Discovery SAGE labs. Animals were singly housed in standard cages with free access to food and water. Study protocols were reviewed and approved by the Stevens Institute of Technology Institutional Animal Care and Use Committee.

### Body weight and genotyping

Rats were weighed weekly using a digital scale. After each rat was sacrificed, the ends of each tail were snipped and sent back to Horizon Discovery SAGE for genotyping to confirm the deletion of the P1 gene.

### Gait analysis

Gait assessment was performed on both the P1KO and age matched WT control rats using a runway system for gait analysis (CSI-G-RWY; CleverSys Inc., Reston, VA). As the rat voluntarily moved across the runway, comprised of a long side-lit glass plate, each footprint was illuminated and recorded by a high speed camera (100 fps) mounted under the glass. Prior to assessment, each rat was trained daily for three consecutive runs until they walked across the apparatus without hesitation. Each rat was then evaluated at 5 months and 8 months of age. Data from three consecutive runs were averaged for each animal. Each video was analyzed using the gait analysis software (GaitScan; CleverSys Inc., Reston, VA).

Analysis was divided into temporal (timing and synchronicity of foot-strike and toe-off events) and spatial measures (geometric characteristics of footprint pattern) of gait (Fig. 1) [20]. Temporal measures (measured in ms) included swing time (time in which paw is in the air), stance time (time in which all paws are detected on the glass) and stride time (stance time plus swing time) (Fig. 1A). Spatial measures (measured in mm) included base of support (average width between hind paws) and stride length (distance the paw traversed from start of previous stance to beginning of next stance) (Fig. 1B-C). An additional measure calculated was the run speed (instantaneous speed over a running distance).

**Figure 1:**
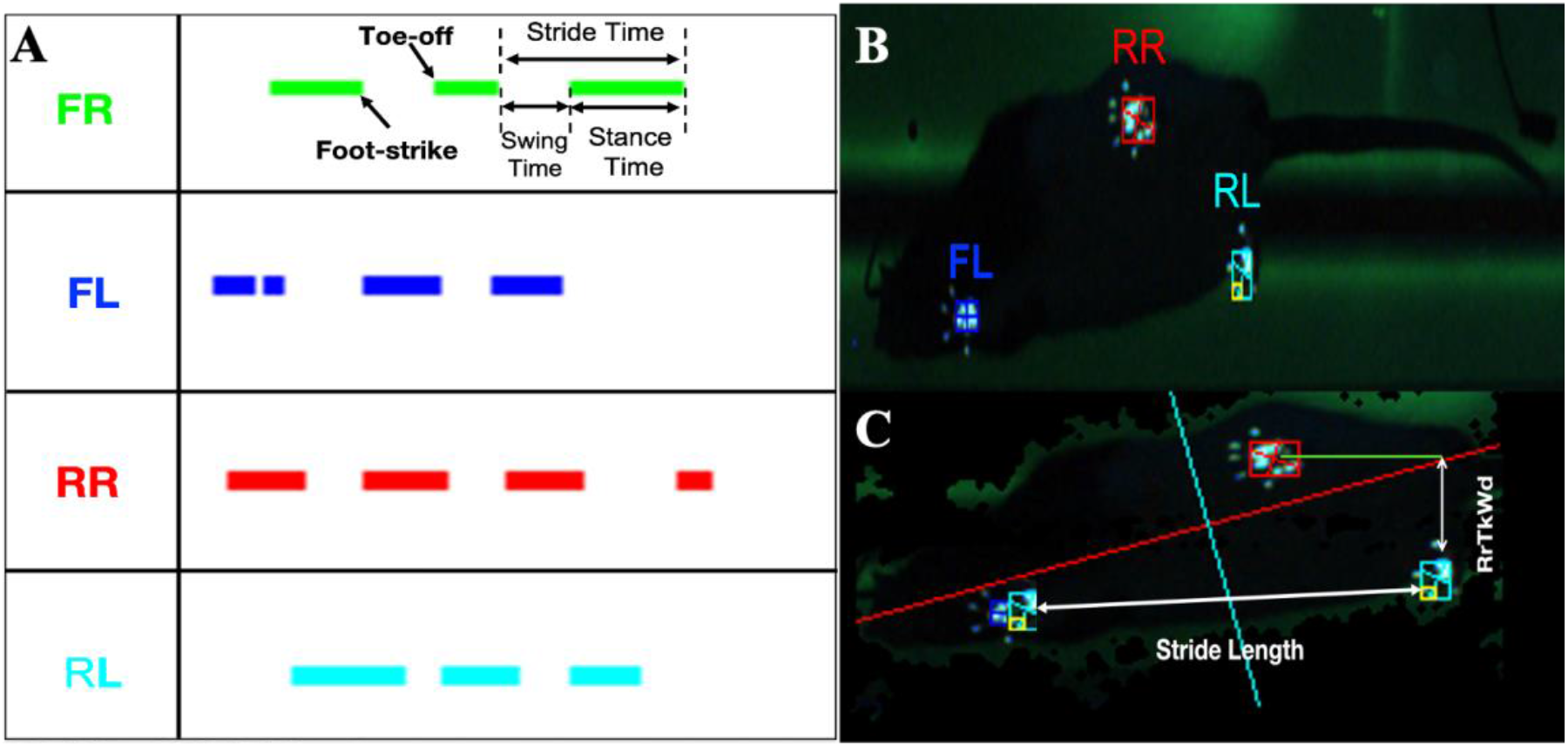
Temporal and spatial characteristics of gait. (A) Temporal gait (measured in ms) include swing time (time in which paw is in the air), stance time (time in which all paws are detected on the glass) and stride time (stance time plus swing time). (B) Foot tracking performed by CleverSys gait scan. (C) Spatial gait (measured in mm) include base of support (BOS, average width between hind paws) and stride length (distance the paw traversed from start of previous stance to beginning of next stance). FR=front right, FL= front left, RR= rear right, RL= rear left.

### Immunohistochemistry

Each animal was deeply anesthetized and transcardially perfused with 1X phosphate buffered saline (PBS) followed by 10% formalin. The brain was removed and postfixed overnight (4°C) in formalin then placed in a 30% sucrose (4°C) solution until it sank. A green tissue marking dye was applied to the left-posterior hemisphere to demarcate the orientation of the brain sections. The brains were cryoprotected with Tissue-Tek optimal cutting temperature (O.C.T.) compound and 40 μm serial coronal sections were cut using a cryostat (CryoStar NX50) equally spaced through the SNc, striatum, and PPN. Immunohistochemistry was performed in the SNc and striatum with anti-tyrosine hydroxylase (TH) antibody to measure dopaminergic neuron loss and on the PPN with anti-choline acetyltransferase (ChAT) antibody to measure cholinergic neuron loss.

#### Anti-TH staining

After three rinses in PBS, sections were blocked for 1 h at 4°C in blocking solution containing PBS, normal goat serum (NGS), and 10% Triton-X. The sections were then incubated in anti-TH monoclonal rabbit IgG primary antibody (1:2000 overnight at 4°C; Sigma-Aldrich) in solution with PBS and NGS. Next, the sections were incubated with goat anti-rabbit IgG Alexa 488 secondary antibody (1:500 for 2 h at 4°C; Invitrogen) in solution with PBS, NGS, and 10% Triton-X. Sections were mounted in DAPI-FluoroMount-G and imaged using a Keyence BZ-X700 series microscope (Fig. 2A-B).

**Figure 2:**
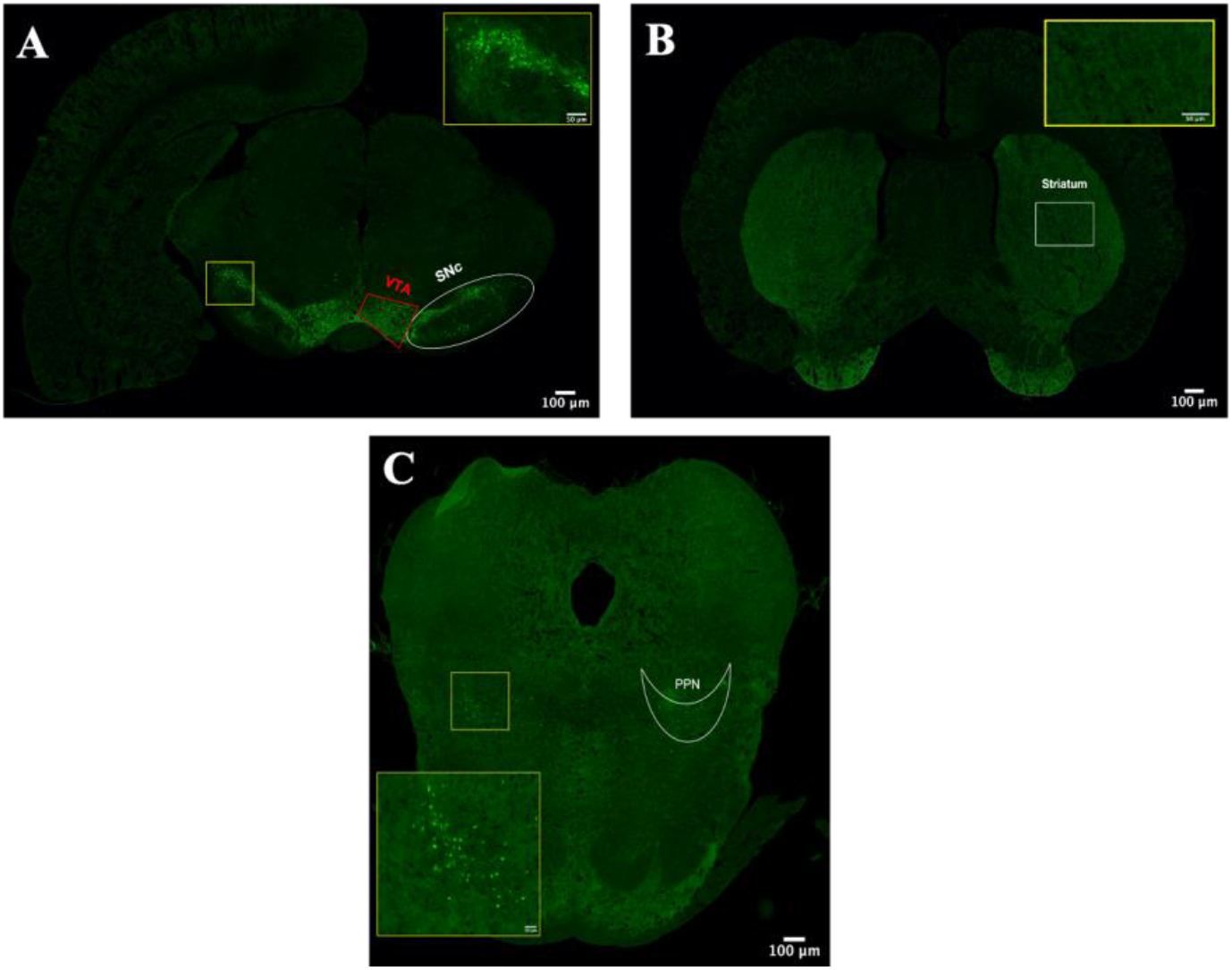
WT sample brain sections. (A) Anti-TH staining to measure dopaminergic neurons in the substantia nigra pars compacta (SNc) and ventral tangential area (VTA) and (B) dopaminergic terminals in the striatum. (C) Anti-ChAT staining to measure cholinergic neurons in the pendunculopontine nucleus (PPN).

#### Anti-ChAT staining

After three rinses in PBS, sections were blocked for 1 h at 4°C in blocking solution containing PBS, normal goat serum (NGS), and 10% Triton-X. The sections were then incubated in anti-ChAT mouse IgG_1_ primary antibody (1:2000 overnight at 4°C; Sigma-Aldrich) in solution with PBS and NGS. Next, the sections were incubated with goat anti-mouse IgG_1_ Alexa 488 secondary antibody (1:500 for 2 h at 4°C; Invitrogen by ThermoFisher Scientific) in solution with PBS, NGS, and 10% Triton-X. Sections were mounted in DAPI-FluoroMount-G and imaged using a Keyence BZ-X700 series microscope (Fig. 2C).

### Cell counting

Automated cell counting and cell fluorescence were performed using ImageJ [21]. Dopaminergic degeneration was determined in the SNc and the VTA by counting the number of TH-positive cells and by measuring cell florescence (pixel intensity) in the striatum of each rat (Fig. 2A-B). Cholinergic degeneration was evaluated in the PPN by ChAT-positive cell counting (Fig. 2C). Cell counts were obtained from a total of five sections spanning each brain region. Automated cell counting was performed bilaterally following the flow chart in Supplementary Fig. 2 [22]. Each 10x image was converted from RGB to 16-bit greyscale. The threshold was set to a pixel intensity of 30 (arbitrary units) and cells were analyzed in a region of interest (ROI) restricted to a 705.9 mm (w) × 564.7 mm (h) rectangle on both sides of the midline in the SNc, 349.0 mm (w) × 458.0 mm (h) in VTA, 352.9 mm (w) × 211.8 mm (h) in striatum, and 493.4 mm (w) × 360.3 mm (h) in PPN (Supplementary Fig. 1B).

### Statistical analysis

Statistical analysis was performed using IBM SPSS Statistics for Mac (IBM Corp., Armonk, N.Y., USA). Means were compared using a two-way mixed ANOVA with age and genotype as factors for gait comparisons between genotypes (P1KO and WT). For significant interactions univariate and one-way repeated measures ANOVA was used to determine significance between genotype at each time point and between timepoints for each genotype, respectively. Variability within the P1KO model was measured by comparing the outcomes of each individual rat across time points. A two-way repeated measures ANOVA was used to compare mean values for each foot for both spatial and temporal measures of gait. Finally, neurotransmitter levels of the P1KO genotype were compared to WT at 8 months of age using a one-way ANOVA. Fisher’s Least Significant Differences (LSD) post hoc test was used for all analyses. Results were considered statistically significant at *p* < 0.05.

## Results

### Bodyweight and genotyping

A two-way mixed measures ANOVA revealed a significant interaction between genotype (P1KO and WT) and behavioral analysis timepoint (5 months and 8 months) (F(1,5)=10.958, *p* = 0.021). Post hoc tests revealed that P1KO rats weighed significantly more at the 5 month test point than the 8 month test point while WT rats remained the same weight between timepoints (P1KO: mean ± SD; WT: mean ± SD). There was no significant difference in bodyweight between genotypes at any given time. Genotyping was confirmed for P1KO rats by gel electrophoresis (Supplementary Fig. 3).

**Figure 3:**
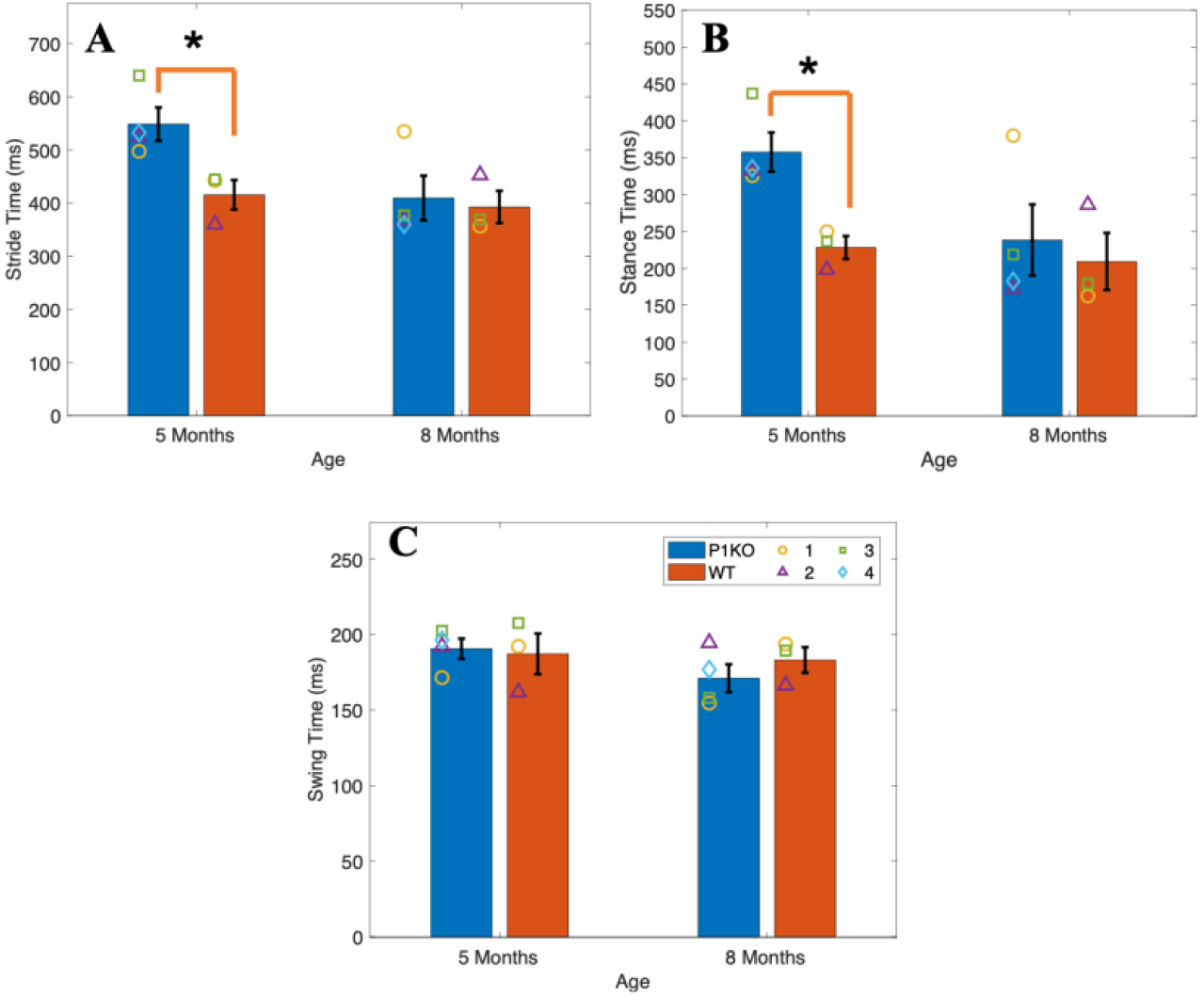
Temporal gait measures in P1KO and WT rats at 5 months and 8 months. (A) Stride time was significantly greater in P1KO rats compared to WT at 5 months but not at 8 months. (B) Stance time was significantly greater in P1KO rats compared to WT at five months but not at 8 months. (C) There were no significant differences in swing time at either time point. Symbols represent the mean at each timepoint in P1KO rats (*n* = 4) and WT control rats (*n* = 3). * indicates *p* < 0.05.

### Gait analysis

Gait dysfunction was observed in both temporal and spatial measures in P1KO rats compared to WT control rats. Visual differences in gait were observed between P1KO and WT controls. Each rat appeared to walk with normal cadence (as seen on the runway gait recordings) except for P1KO3 whose hindlimbs dragged and swung forward at the same time during ambulation, mimicking a hopping motion (Supplementary Fig. 4). This anomalous gait resulted in an increase in double-limb support time (data not shown) compared to the remaining P1KO rats.

**Figure 4:**
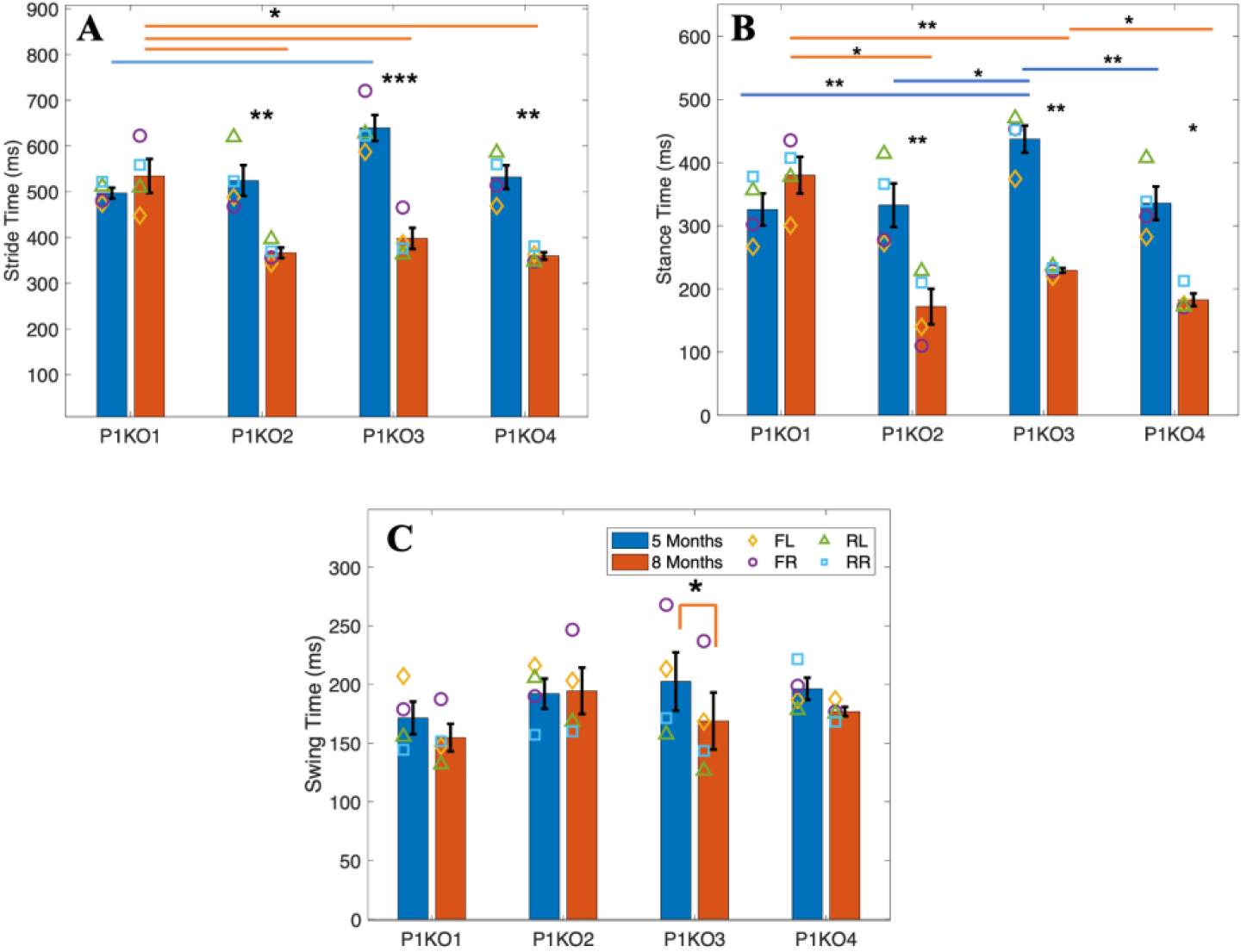
Temporal gait measures in P1KO rats at 5 months and 8 months. (A) P1KO3 had the greatest stride time at 5 months. There was a significant difference in stride time between P1KO1 and P1KO3 at 5 months and P1KO1 and P1KO2, P1KO3, and P1KO4 at 8 months. Each rat except for P1KO1 experienced a significant improvement in stride time from 5 to 8 months. (B) P1KO3 had the longest stance time at 5 months which was significantly greater than all other P1KO rats. There was a significant improvement in gait for all rats at 8 months, except for P1KO1 who experienced an increase in stance time. (C) There were no significant differences in swing time between animals, however, P1KO3 displayed a significant decrease in stance time over time. Symbols represent the mean temporal measure of each paw. * indicates *p* < 0.05; ** indicates *p* < 0.01; *** indicates *p* < 0.001.

#### Temporal measures

Stride time, stance time, and swing time were compared between P1KO rats and WT controls. P1KO rats experienced gait deficits compared to age matched WT controls only at 5 months of age. No statistically significant interaction between genotype and time was found for stance time, swing time, and stride time. Significant differences were found for the main effect genotype stride time (F(1,5) = 10.755, *p* = 0.022) but not for stance time (F(1,5) = 6.460, *p* = 0.052) or swing time (F(1,5) = 0.137, *p* = 0.726). P1KO had a significantly higher stride time compared to the WT control. Simple effects analysis revealed significant differences between P1KO and WT at 5 months but not at 8 months for both stride time (F(1,5) =9.215 p = 0.029) and stance time F(1,5) = 14.481 *p* = 0.013). No significant main effect of time was found across temporal gait measures. Overall, stance time and stride time were greater in P1KO rats than WT at 5 months but had similar means at 8 months (Fig. 3A-B). Swing time in the P1KO model was comparable to WT at both time points (Fig. 3C).

Individual analysis within the P1KO model revealed a significant interaction between P1KO rat and time for stride time (F(3,9) = 27.133, *p* < 0.001) and stance time (F(3,9) = 38.803, *p* < 0.001) (Fig. 4A-B). No significant interaction was found for swing time (Fig. 4C).

Simple main effects on stride time for rat found significance between P1KO rats at 5 months (F(3,9)=7.356, *p* = 0.009) and 8 months (F(3,9) = 14.414, *p* = 0.001).

Post hoc analysis found significance between P1KO1 and P1KO3 (*p* = 0.022) at 5 months and between P1KO1 and P1KO2 (*p* = 0.021), P1KO3 (*p* = 0.014), and P1KO4 (*p* = 0.021) at 8 months. P1KO1 had the lowest mean stride time at 5 months and the highest at 8 months compared to the other test subjects. Simple main effects for time revealed a significant decrease in stride time for P1KO2 (F(1,3) = 45.521, *p* = 0.007), P1KO3 (F(1,3) = 281.711, *p* < 0.001), and P1KO4 (F(1,3) = 1114.165, *p* < 0.001) from 5 months to 8 months (Fig. 4A).

Simple main effects for stance time revealed significant differences between P1KO rats overall at 5 months (F(3,9) = 16.870, *p* < 0.001) and at 8 months (F(3,9) = 24.258, *p* < 0.001). Post hoc analysis showed that stance time for P1KO3 was significantly longer than P1KO1 (*p* = 0.035) and P1KO4 (*p* = 0.47) at 5 months. P1KO1’s stance time was significantly longer than P1KO4 (*p* =0.038) at 8 months

Simple main effects also found significant differences in stance time between 5 months and 8 months in P1KO2 (F(1,3) = 205.923, *p* = 0.001), P1KO3 (F(1,3) = 134.530), and P1KO4 (F(1,3) = 28.794, *p* = 0.013). Stance time at 5 months was greater than stance time at 8 months in each rat where significance was found (Fig. 4B).

Although there were no significant interactions or main effects for swing time between P1KO and WT or between P1KO rats, there was a significant asymmetry in P1KO3 rat at 5 months (F(3,6) = 8.687, *p* = 0.013). Post hoc analysis revealed significant differences between the front left and front right paw (*p* = 0.040) and rear right and rear left (*p* = 0.014) (Fig. 4C). This asymmetry resolved itself at 8 months. No significance in swing time asymmetry was present in the other P1KO rats.

#### Spatial measures

There was no significant two-way interaction between genotype and time for both stride length and base of support (Fig. 5). Main effect of genotype was not significant for either measure, while main effect of time showed a statistically significant difference in stride length ((F(1,5) = 23.479, *p* = 0.005) between time points. Post hoc analysis revealed a significant decrease in stride length from 5 to 8 months in P1KO rats (F(1,3) = 15.930, *p* = 0.028), but not WT (Fig. 5B).

**Figure 5:**
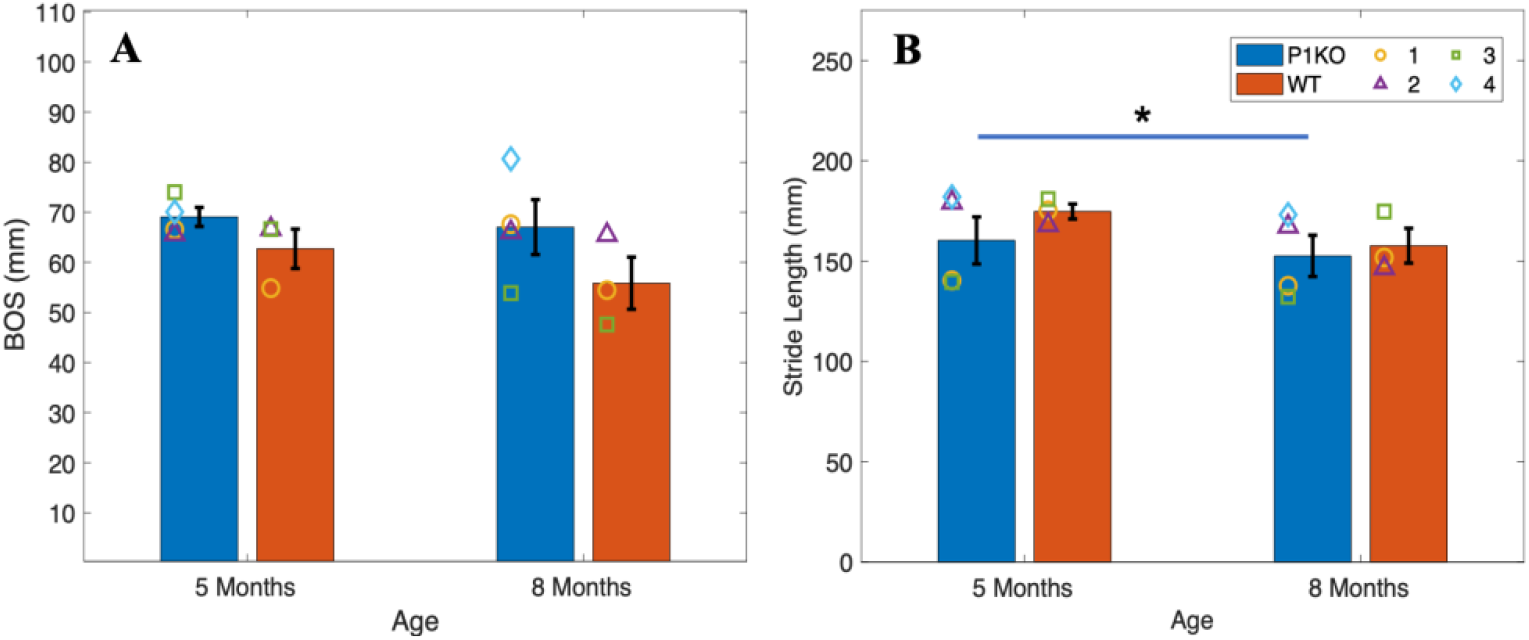
Spatial gait measures in P1KO and WT rats at 5 months and 8 months. (A) Rear track width (RrTkWd) was similar across genotype and time. (B) Stride length remained constant between genotypes at each time point. There was a significant decrease in P1KO stride length from 5 to 8 months. Symbols represent the mean at each timepoint in *n* = 4 P1KO rats and *n* = 3 WT control rats. * indicates *p* < 0.05.

No significant interaction between P1KO rat and time was found for stride length or base of support (Fig. 6). Main effect of differences between knockout rats over time was not significant for either measure, while main effect of time showed a statistically significant difference in stride length (F(1,3) = 15.930, *p* < 0.01) between time points. Post hoc analysis found significant differences in stride length between P1KO1 and P1KO3 (*p* = 0.008), P1KO1 and P1KO4 (*p* < 0.01), P1KO2 and P1KO3 (*p* = 0.047), and P1KO3 and P1KO4 (*p* = 0.011) over time (Fig. 6B).

**Figure 6:**
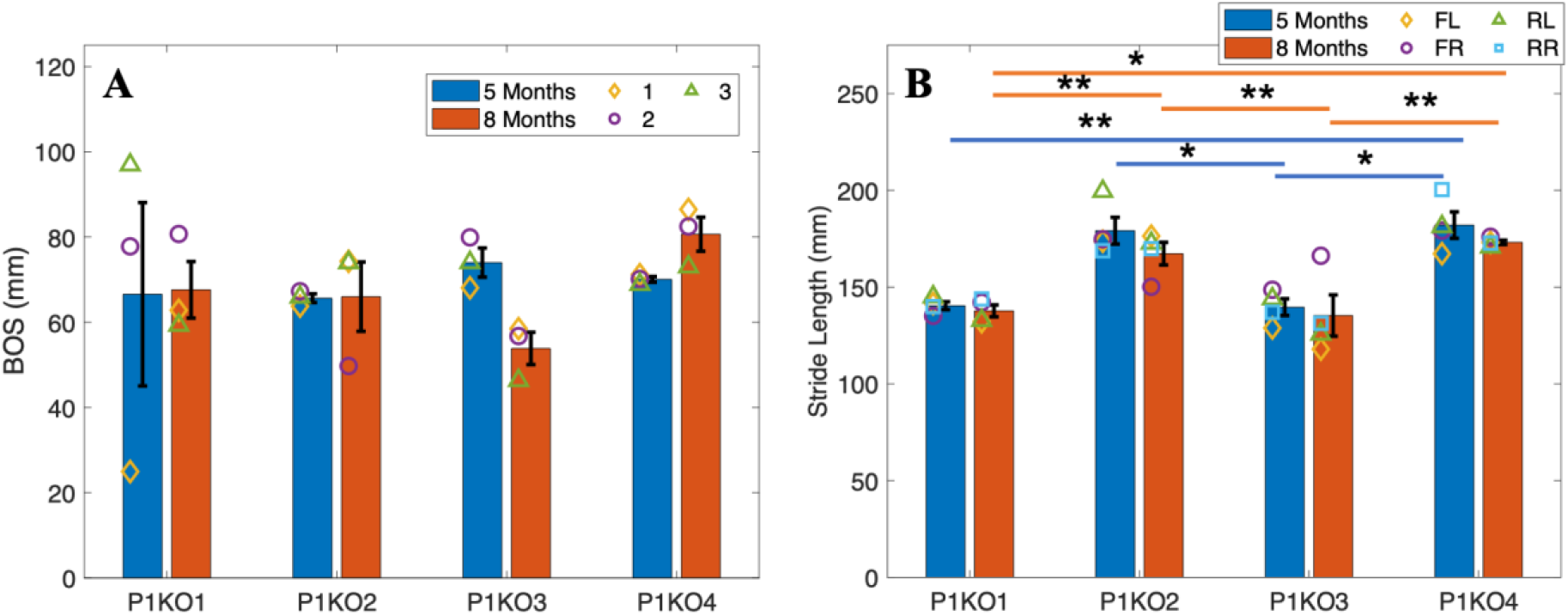
Spatial gait measures in P1KO rats at 5 months and 8 months. (A) There were no significant differences in base of support (BOS) between rats or over time. P1KO3 experienced a decrease in BOS while each other rat stayed the same or had a slight increase. (B) Significant differences were found in stride length between rats at both 5 months and 8 months. A significant difference between P1KO3 and P1KO2 and P1KO4 was measured at 5 months and between P1KO1 and P1KO2, P1KO2 and P1KO3, P1KO3 and P1KO4, and P1KO1 and P1KO4 at 8 months. There were no significant changes in gait between 5 months and 8 months. Symbols represent the mean temporal measure of each paw. * indicates *p* < 0.05; ** indicates *p* < 0.01; *** indicates *p* < 0.001.

Simple main effects for stride length revealed significant differences between P1KO rats overall at 5 months (F(3,9) = 20.398, *p* < 0.001) and at 8 months (F(3,9) = 8.129, *p* = 0.006) (Fig. 6B). At 5 months, stride length for: 1) P1KO1 was significantly shorter than P1KO2 (*p* = 0.007) and P1KO4 (*p* = 0.011); 2) P1KO2 was significantly larger than P1KO3 (*p* = 0.009); 3) P1KO3 was significantly shorter than P1KO4 (*p* = 0.010). At 8 months, stride length: 1) for P1KO1 was significantly shorter than P1KO2 (*p* = 0.036) and P1KO4 (*p* = 0.001); 2) and for P1KO3 was significantly shorter than P1KO4 (*p* = 0.031). Overall, stride length at 5 months was greater than at 8 months for each animal (Fig. 6B).

#### Run speed

There was no significant two-way interactions between genotype and time on run speed. The main effect for genotype was statistically significant (F(1,5)=18.334, *p* = 0.008). Simple effects showed a significant difference between P1KO and WT rats at 5 months (*p* < 0.01) and 8 months (*p* < 0.05). P1KO rats had a slower instantaneous speed than WT at each time point (Fig. 7A).

**Figure 7:**
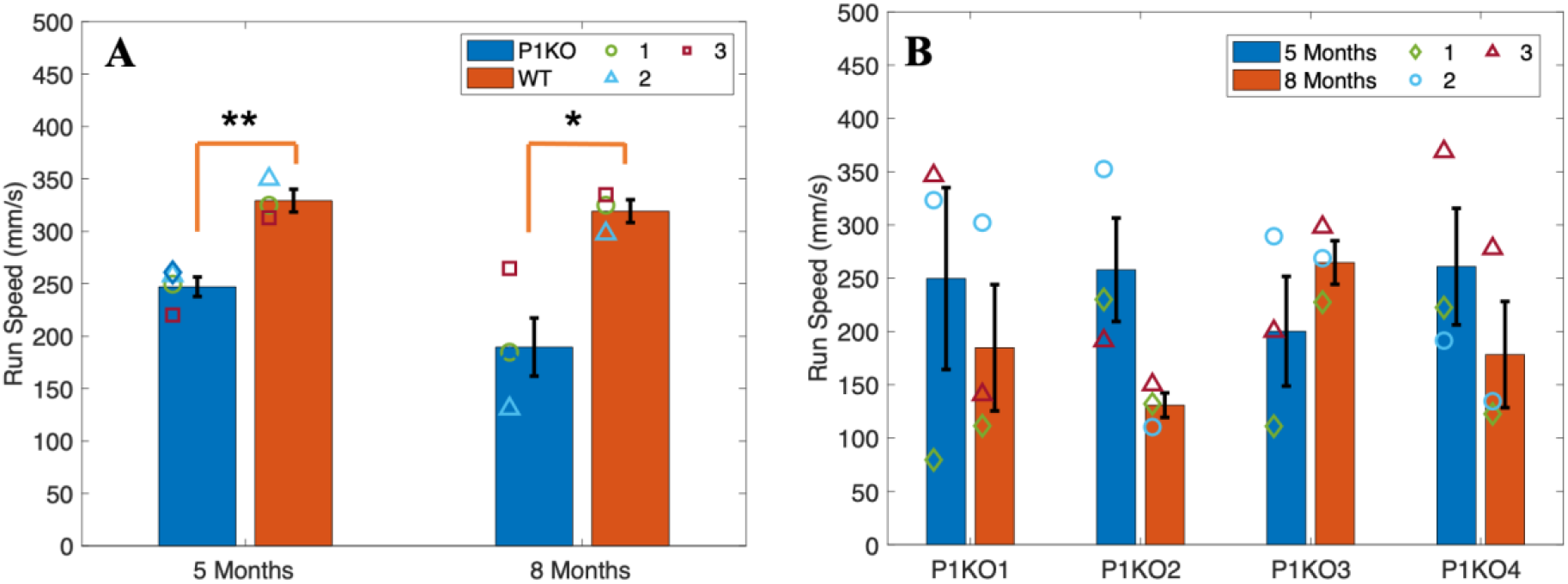
Run speed. (A) P1KO had a significantly slower run speed than WT controls at both 5 months and 8 months. P1KO moved more slowly over time (although not significantly) while WT’s speed remained the same. Symbols represent the mean at each timepoint in *n* = 4 P1KO rats and *n* = 3 WT control rats. (B) Individual comparison between P1KO rats. There were no significant differences in run speed observed between rats over time. Symbols represent the mean temporal measure of each paw. * indicates *p* < 0.05; ** indicates *p* < 0.01.

No significant interaction was found between P1KO rat and time or for main effects of genotype and time. P1KO1 and P1KO3 were faster than P1KO2 and P1KO4 at 8 months (Fig. 7B).

### Cell Counting

Cell counting in the SNc resulted in a 26.7% loss of TH+ cells in P1KO compared to WT (F(1,6) = 8.075, *p* = 0.030). There was no significant difference in SNc TH+ cells between P1KO rats (Fig 8A). Fig. 8B compares the number of TH+ cells in the SNc of each P1KO rat to both temporal and spatial gait parameters. Mean cell counts in order from greatest to least are P1KO2 (472.667), P1KO1 (418.333), P1KO4 (391.250), and P1KO3 (338.833). The number of TH+ cells in the VTA was not reduced in P1KO rats (Fig. 8C). A 15% loss of TH+ terminals was observed in the striatum (F(1,6) = 6.035, *p* = 0.049) (Fig. 8D). No differences were observed in ChAT+ cells between P1KO and WT in the PPN (Fig. 8E).

**Figure 8:**
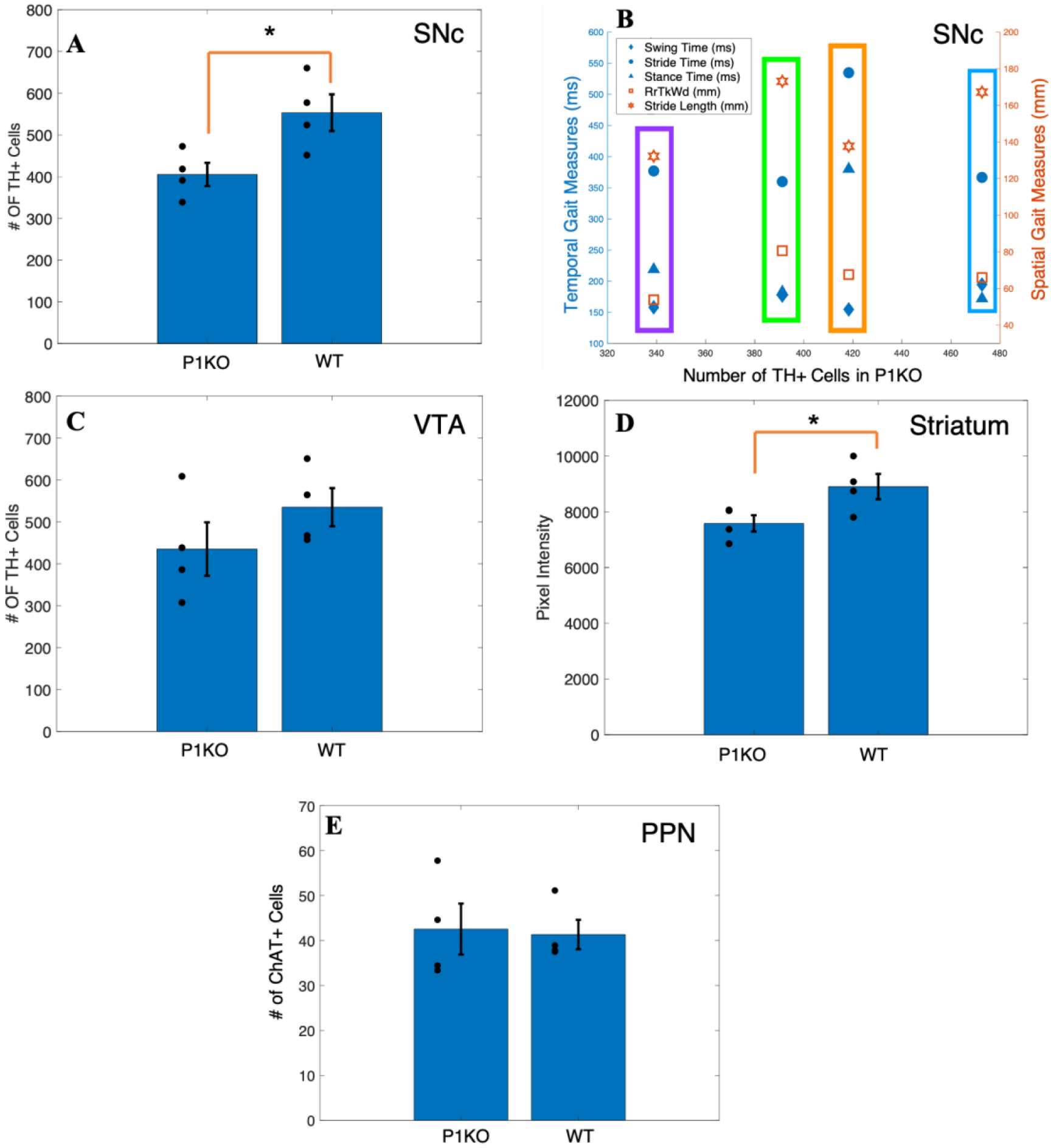
Cell counts. (A) P1KO had significantly less TH+ cells in the substantia nigra pars compacta (SNc) than WT at 8 months. (B) Comparison between spatial and temporal gait deficits and TH+ cell counts in the SNc of each P1KO rat at 8 months. The boxes represent individual P1KO rats while each symbol represents spatial and temporal parameters of gait. Purple = P1KO3, green = P1KO4, orange = P1KO1, and blue= P1KO1. (C) No significant difference in the number of TH+ cells in the VTA between genotypes. (D) There was a significant decrease in the expression of TH+ terminals in the striatum by P1KO and WT rats as measured by the average pixel intensity over region of interest. (E) There was no significant difference in the number of ChAT+ cells in the pendunculopontine nucleus (PPN) between genotypes. Each point represents the mean cell count for P1KO (*n* = 4) rats and WT (*n* = 4) rats. * indicates *p* < 0.05; ** indicates *p* < 0.01.

## Discussion

Parkinson’s disease (PD) is a progressive neurodegenerative movement disorder that causes gait dysfunction typically manifesting in the later stages of the disease. Clinically, symptoms present as a decrease in stride length, velocity (run speed), and swing time, increase in stride time and stance time, and an increased base of support [20,23,24,25]. An age dependent and progressive animal model of PD is needed to assist in finding effective treatments for patients experiencing these gait deficits [8]. The P1KO rat model of PD may fill this need, however, many studies have not directly tested gait parameters that relate to what is observed clinically and those that have, produced varying results. To determine if the P1KO is a consistent and feasible model of progressive gait degeneration in PD, gait analysis was performed in male P1KO and WT control age matched rats at 5 months and 8 months of age.

Gait disturbances measured at 5 months parallel clinical observations, although not all differences reached statistical significance. Five months aged P1KO rats experienced a significant increase in stance time and stride time and a significant decrease in run speed compared to age matched WT controls. Rats lacking the P1 gene moved more slowly and spent more time with their paws on the runway, mimicking typical symptoms of PD. P1KO stride length, swing time, and base of support were not significantly different than WT. These findings support the face validity of the P1KO rat model for gait dysfunction in PD.

At 8 months of age, spatial and temporal gait measurements in P1KO rats more closely resembled those found in WT, while WT remained unchanged over time. P1KO mean stance time, stride time, swing time, stride length, and run speed decreased and base of support stayed the same. These changes, however, were not significant compared to the same measures at 5 months. Overall, gait disturbances initially present at 5 months seemed to improve by 8 months. Similar findings were reported in 8 month old female P1KO rats suggesting this may be attributed to compensatory adjustments occurring during behavioral testing [26]. Dave *et al.* [13] and de Haas *et al.* [11], however, reported significant gait deficits continuing up to 8 months of age. Dave *et al.* [13] found a significant decrease in stride time, stance time, and swing time compared to age matched WT controls at 8 months using the neural cube behavioral apparatus. These findings are atypical of what is observed clinically in PD and in this study. de Haas *et al*. [11] reported that at 8 months the P1KO model exhibited gait deficits (stance time and swing time) similar to our results at 5 months. In addition, de Haas *et al.* [11] found a significant increase in base of support and a significant decrease in stride length. They did not perform behavioral analysis at any other time point.

Variability between studies indicates variability within the P1KO model itself. Dave *at al*. found that in one subset of rats, six out of fifteen P1KO rats exhibited impaired gait at 8 months and fifteen out of fifteen P1KO rats exhibited normal gait at 4 and 6 months [13]. In the second subset of 7 P1KO rats tested, all rats showed gait dysfunction at 4 months. These results suggest that not all rats will experience gait dysfunction, but those that do tend to present symptoms between 4 and 8 months of age. All P1KO rats in the current study showed gait deficits with varying severity at 5 months. At 5 months, one out of four P1KO rats (P1KO3) exhibited the most severe gait dysfunction across all measures. This was the only rat with gait deficits that could be readily observed as it ran across the runway. The rat would lift and drag its hindlegs together as it walked (Supplementary Fig. 5). At 5 months, P1KO3 had the greatest stride time, stance time, and base of support and the lowest run speed and stride length (Fig. 4, 6, 7). This was the only subject to exhibit asymmetry in swing time which is directly correlated to freezing of gait [27].

Stride length and stance time showed a different trend compared to other P1KO symptoms. At 8 months, stride length stayed the same in all rats while stride time and stance time decreased significantly in three out of four rats. Dragging persisted in P1KO3 at 8 months, but the rat showed some compensation for this deficit, as shown by improvement in gait parameters. Hind limb dragging was also observed by Dave *et al*. [13] at 6 months but resolved by 8 months. P1KO1 was the only rat to experience worsened gait from 5 months to 8 months including increased stride length and stance time, and decreased swing time and run speed.

A pathological hallmark of PD is dopamine degeneration in the SNc and striatum. Gait dysfunction in the P1KO model may be attributed to DA loss in the SNc and striatum. We measured a 27% loss of TH+ cells in the SNc and 15% loss of TH+ terminals in the striatum in P1KO aged 8 months compared to WT controls. The same P1KO rat with the greatest gait deficits showed the greatest loss of TH+ cells in the SNc.

Clinically, motor symptoms appear in more advanced stages of PD when 60% to 80% of striatal dopaminergic terminals and 50% of dopaminergic neurons in the SNc are lost [28, 29, 30]. Generally, this does not seem to be the case in the P1KO model. To our knowledge, the current study is the first to report a significant decrease in the intensity of TH+ terminals in the striatum in addition to significant loss of TH+ cells in the SNc. Most rats, however, present motor symptoms either prior to DA loss in the SNc or without exhibiting DA loss in the SNc [11,13,14,15,26]. Select studies measuring DA loss found that DA loss in the SNc coincided with an increase in striatal DA [13,16].

Although loss of DA in the striatum and SNc has not been consistently reported, nigrostriatal damage may underlie gait dysfunction in P1KO rats. An increase in striatal DA may be a hallmark for the P1KO rat model and predictive of the onset of DA loss in the SNc. Metabolic alterations in the P1KO striatum occur prior to loss of dopaminergic neurons in the SNc at 9 months of age [17]. Creed *et al.* [16] observed an increase in striatal DA at 8 months, which eventually decreased to a point of significant loss at 12 months. It has been suggested that the increase in dopaminergic terminals in the striatum is a compensatory mechanism for early nigrostriatal damage that disappears as the disease progresses [31,32]. There exists a paucity of studies that measured dopamine at multiple time points in P1KO rats. Like the current study, most studies performed histochemistry on the P1KO striatum and SNc only at 8 months of age [11,13, 14,15,26], however, changes that occurred in striatal DA at either earlier or later time points could not be determined. In the current study, histochemistry at 8 months *per se* allowed for longitudinal gait measurements within the same rats at both the 5 month and 8 month time points. A limitation of this experimental design was the inability to analyze TH expression at 5 months of age in the same animals. Future studies are needed to quantify DA loss in the SNc and striatum from 2 months (prior to the onset of motor abnormalities) through at least 12 months across a large number of P1KO and WT rats.

Another possible cause of further gait degeneration displayed by the P1KO model is a loss of cholinergic neurons in the PPN. Loss of cholinergic neurons in the PPN is believed to be a cause of gait and balance deficits in PD patients [4]. There was no significant loss of ChAT+ cells in the PPN suggesting that gait disturbances in PINK1 KO rats are not due to PPN degeneration.

The results presented here and those reported by previous studies highlight the variability of the P1KO model. Most studies analyzed P1KO rats bred by SAGE labs [11,12,13,14,15]. Two out of three studies that reported neurodegeneration of DA used P1KO rats from an undisclosed source [16,17]. These rats may have been bred in house. Housing conditions, stress, microbiome, and food can all play a role in neurodegeneration [7]. Breeding conditions and stress experienced by rats during shipping may account for the presence or lack of neurodegeneration in the P1KO model at 8 months.

Variability within a model of gait dysfunction in PD can be beneficial, because symptoms do not present in the same way or with the same severity in each person. Variation in dysfunction would allow for more realistic results when testing for treatments used to improve Parkinsonian gait. The cause for these gait deficits, however, is unknown. Although most P1KO rats will display gait deficits at some point in their lifetime, these deficits may or may not be attributed to loss of DA in either the SNc or the striatum. Gait treatments targeting the basal ganglia may not give proper insight into how it will affect Parkinsonian patients if one of the hallmark features of PD does not occur. Another factor to consider is that it may take up to 8 months before the rats present with motor symptoms [13], making the studies more expensive due to the prolonged care and maintenance of the rats required. The demonstration of consistent and progressive gait and nigrostriatal degeneration by future studies is necessary for the P1KO model to realize its full potential as an invaluable tool for PD research.

## Funding

This study was supported by a grant from the Branfman Family Foundation (G.C.M.).

## Competing Interests

The authors report no competing interests.

## Abbreviations

DA: dopamine
P1KO: PTEN-induced putative kinase 1 knockout
PBS: phosphate buffered saline
PD: Parkinson’s disease
PPN: pedunculopontine nucleus
SNc: substantia nigra pars compacta
VTA: ventral tegmental area
WT: wild-type

## Notes

### Competing Interest Statement

The authors have declared no competing interest.

