## Supplemental Materials for "Dopaminergic but not Cholinergic Neurodegeneration is Correlated with Gait Disturbances in PINK1 Knockout Rats"

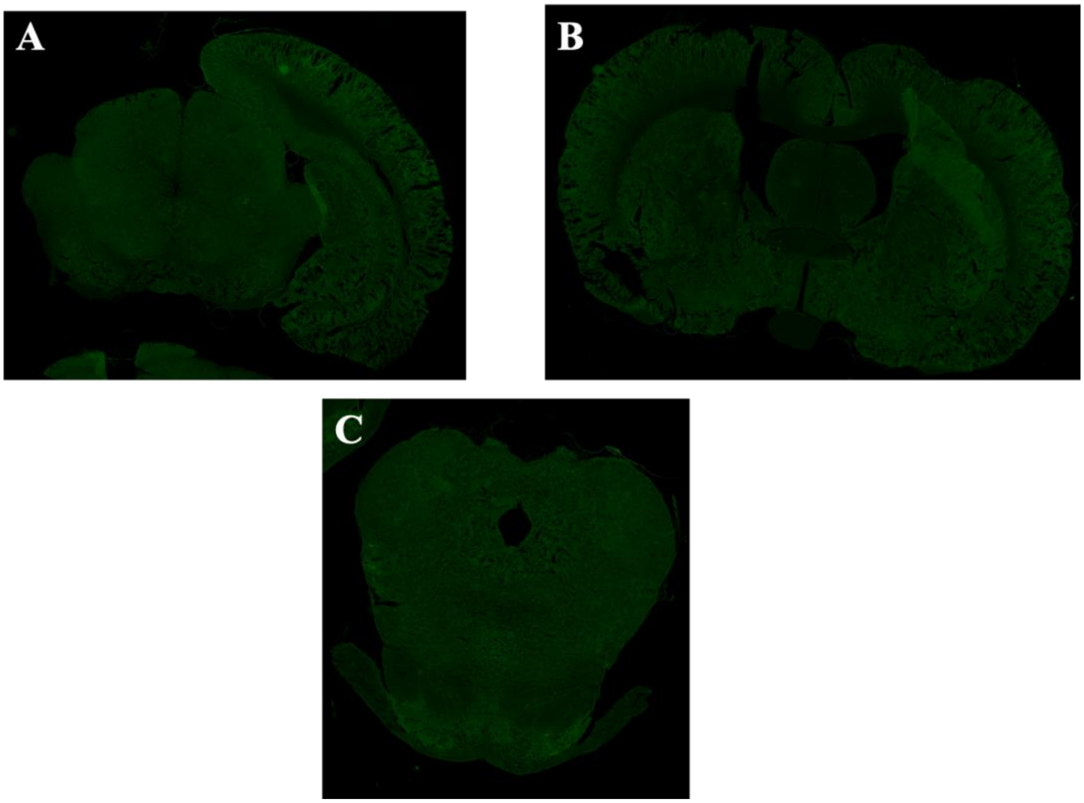


**Supplementary Figure 1: Representative brain sections from the A) SNc B) Striatum and C) PPN stained without addition of primary antibody. SNc = substantia nigra pars compacta; PPN = pedunculopontine nucleus.**

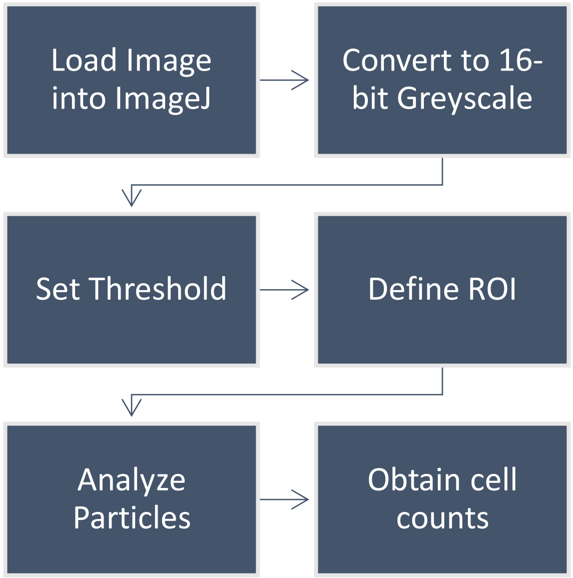


**Supplementary Figure 2: Flowchart for cell counting using ImageJ’s Particle Analyzer.**

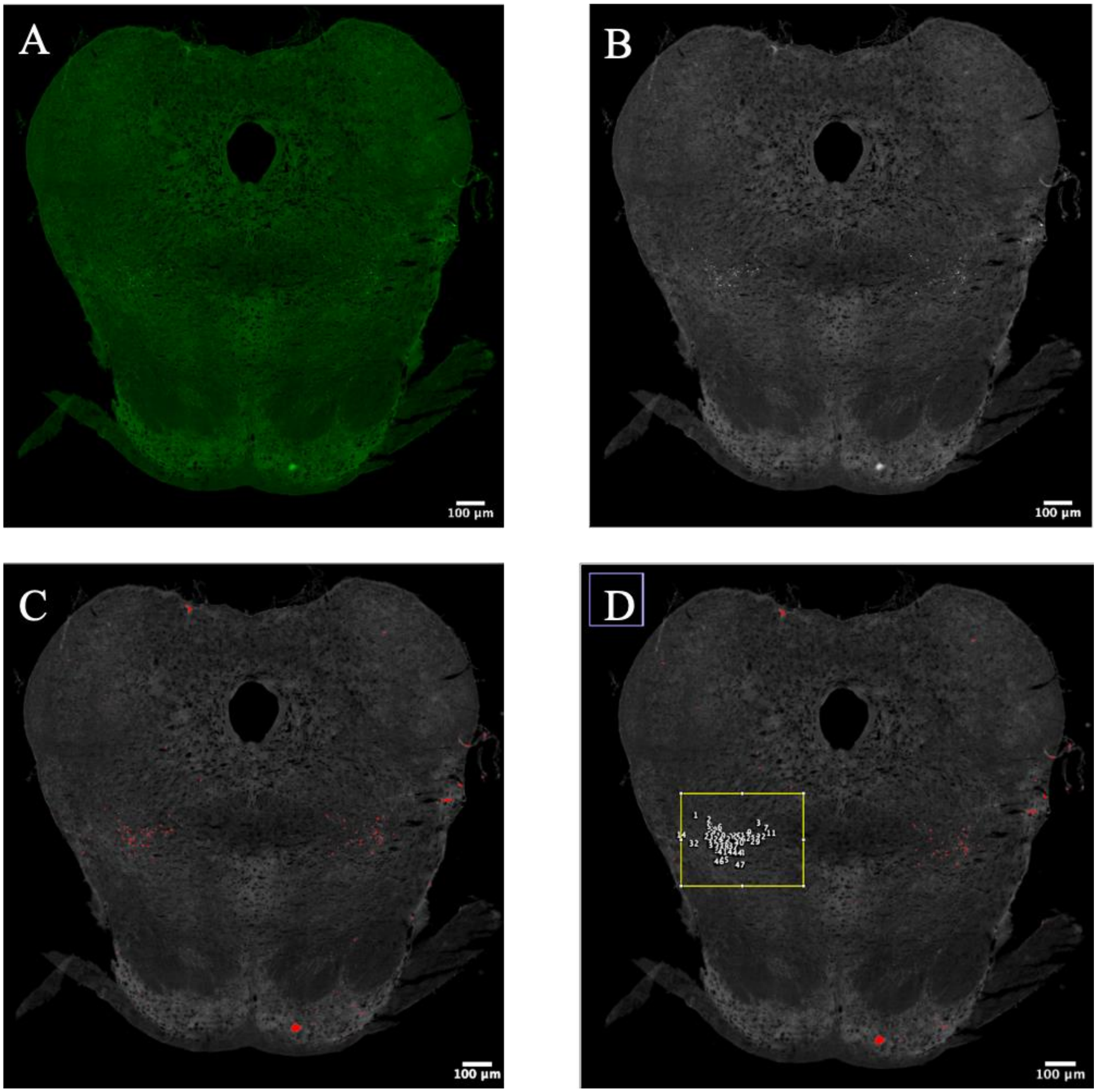


**Supplementary Figure 3: Steps outlining cell counting procedure.** (A) Original image. (B) First, image is converted to a 16-bit greyscale image. (C) Next, the threshold is set to highlight the cells and differentiate from the background. (D) Finally, automatic cell counting is used within a defined area (indicated by yellow box).

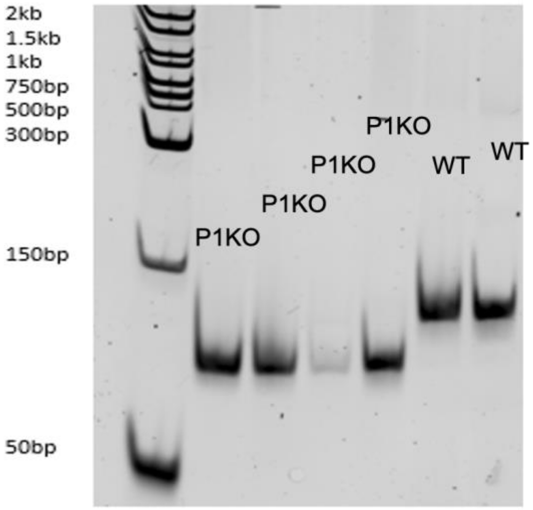


**Supplementary Figure 3: Genotyping results for all four P1KO rats and two of four WT rats used in this study.**

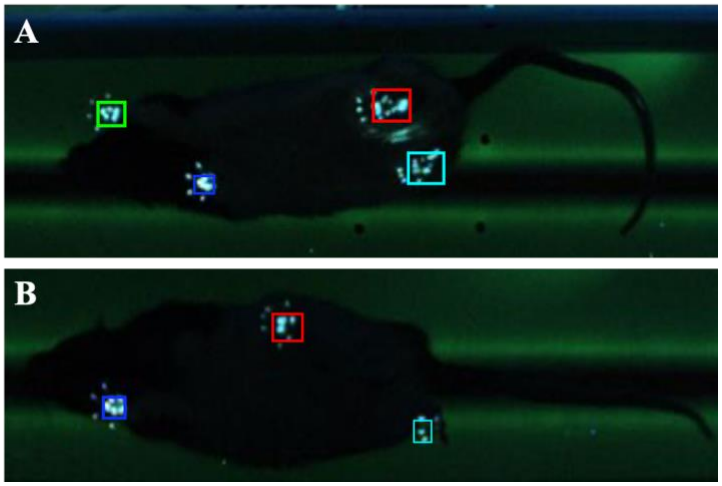


**Supplementary Figure 4: Examples of P1KO and WT stepping pattern.** (A) P1KO3 rat with irregular stepping pattern, rear track width is decreased, and rat lifts or drags both hindlimbs at same time. (B) WT control with normal stepping pattern. Green box indicates the front right paw, dark blue indicates front left paw, red indicates rear right paw, and turquoise indicates rear left paw.

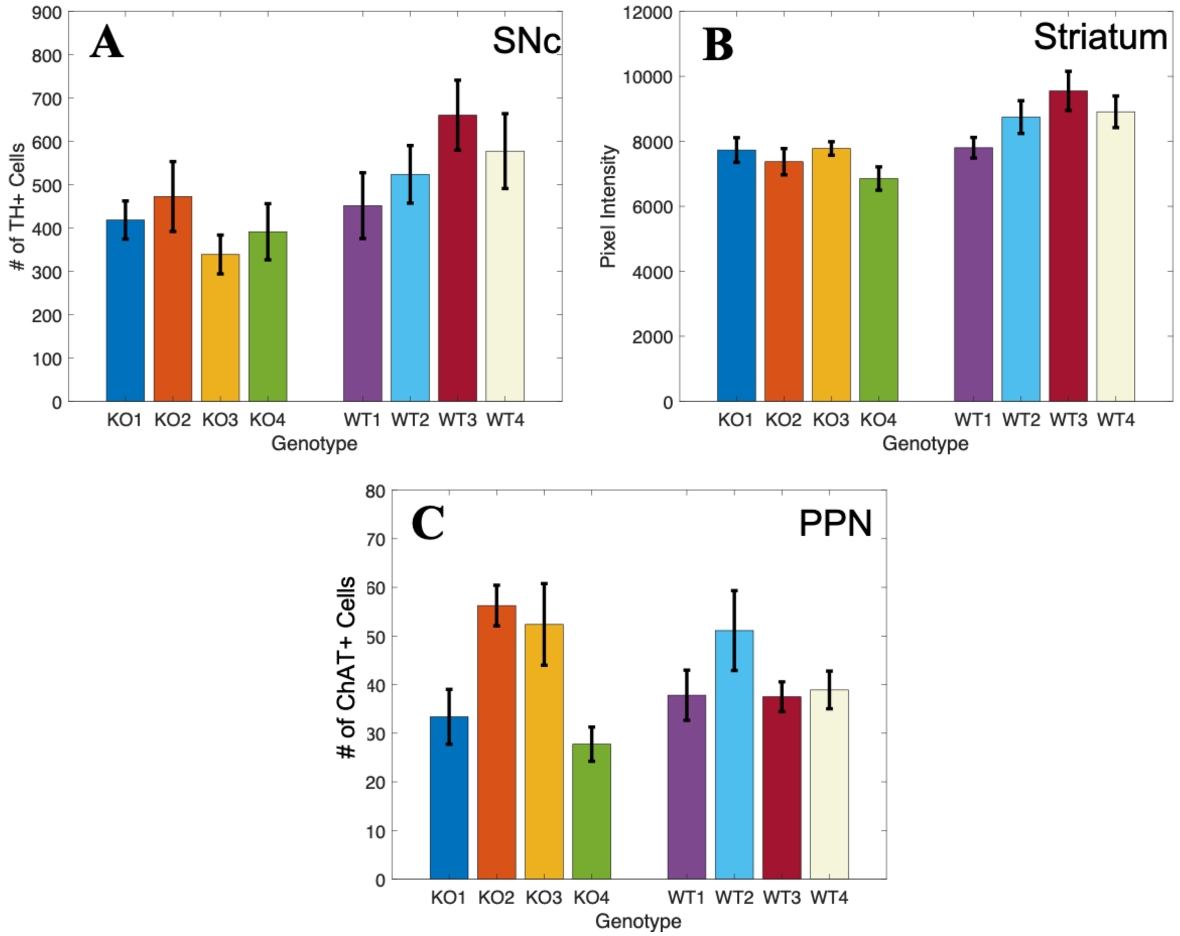


**Supplementary Figure 5: Individual mean cell counts from brain sections of the (A) SNc (*n* = 6), (B) Striatum (*n* = 6), and (C) PPN (*n* = 6).** Abbreviations: SNc = substantia nigra pars compacta; PPN = pendunculopontine nucleus.

**Supplementary Video 1: Gait pattern of a sample wild type rat.** The video speed was reduced to better display the gait pattern.

**Supplementary Video 2: Gait pattern of a sample Pink1KO rat.** The video speed was reduced to better display the gait pattern.
